# *Staphylococcus aureus* Biofilm Removal by Targeting Biofilm-Associated Extracellular Proteins

**DOI:** 10.1101/102384

**Authors:** Sudhir K. Shukla, T. Subba Rao

## Abstract

**Aim:** Among cell surface proteins, biofilm-associated protein promotes biofilm development in *Staphylococcus aureus* strains. Aim of this study was to investigate proteinase-mediated biofilm dispersion in different isolates of *S. aureus*.

**Methods and Results:** Microtitre plate based biofilm assay showed that 2 μg/mL proteinase K significantly inhibited biofilm development in *bap*-positive *S. aureus* V329 as well as other *S. aureus* strains, i.e. SA7, SA10, SA33, SA352 and but not in *bap*-mutant M556 and SA392 (a weak biofilm producing strain). However, proteinase K treatment on *S. aureus* planktonic cells showed that there was no inhibition of planktonic growth at any concentration of proteinase K when tested up to 32 μg/mL. This observation ruled out the possibility of *S. aureus* biofilm inhibition by altering the cell viability. Proteinase K treatment upon 24 h old preformed biofilms showed an enhanced dispersion of *bap*-positive V329 and SA7, SA10, SA33 and SA352 biofilms, however, proteinase K did not affect the *bap*-mutant *S. aureus* M556 and SA392 biofilms. Biofilm compositions study before and after proteinase K treatment indicated that Bap might also be involved in eDNA retention in the biofilm matrix that aid in biofilm stability. When proteinase K was used in combination with antibiotics, a synergistic effect in antibiotic efficacy was observed against all biofilm forming *S. aureus* strains.

**Conclusion:** Proteinase K inhibited biofilms growth in *S. aureus* bovine mastitis isolates but did not affect their planktonic growth. An enhanced dispersion of preformed *S. aureus* biofilms was observed upon proteinase K treatment. Proteinase K treatment with antibiotics showed a synergistic effect against *S. aureus* biofilms.

**Significance of the study:** The study suggests that dispersing *S. aureus* by protease can be of use while devising strategies against *S. aureus* biofilms. Proteinase K treatment has a wider scope for control of *S. aureus* biofilms.

## Introduction

Biofilms are complex microbial communities adhered to biotic or abiotic surfaces and are embedded in a matrix of extracellular polymeric substance (EPS)^1^. Most bacteria in nature execute definite development stages in biofilm formation; a) adherence of cells to a substratum, b) development of micro-colonies, c) maturation of micro-colonies into biofilms and d) detachment of bacteria and acquisition of motile phase, known as biofilm dispersal^2^. Biofilm disassembly/dispersion is believed to play very important role in pathogenicity, environmental distribution and also in phase transition^1, 3^. The dispersal phenomenon can also be triggered by several environmental signals or unfavourable condition^4^. Biofilms aid many advantages to microorganisms such as higher resistance to adverse environmental conditions, higher resistance to antimicrobial agents and enhanced protection from immune response in the case of persistent infections^5^. Therefore, in recent times researchers have targeted biofilm dispersal phenomenon, particularly to enhance the microbial susceptibility towards antimicrobials. Since biofilms are known to be a source of persistent infection, new approaches are needed to combat biofilm mediated persistent infections^6^.

*S. aureus* is a universal pathogen which causes mild to severely life threatening diseases^7^. This bacterium also constitutes a major cause of hospital-acquired/healthcare-associated infections (HAIs). According to Center for Disease Control and prevention (CDC), *S. aureus* strains are associated with 15.6 % of the total HAIs reported between 2009 and 2010,^8^ and 12.3 % between 2011 and 2012 in Europe^9^. Commonly, a mature biofilm consists of polysaccharides, proteins, extracellular DNA^10^ and amyloid fibres as matrix^1^. There are several reports on the role of surface proteins in *S. aureus* biofilm formation and its stability^11-14^. Among various surface proteins, biofilm-associated-protein (Bap) was first reported as a large, multi-domain, cell surface anchored protein, which plays a crucial role in *S. aureus* biofilm development, architecture and in the pathogenesis of bovine mastitis^15-19^. A recent study carried out in Brazil showed the presence of *bap* gene in all the coagulase-negative *Staphylococcus spp.* strains isolated from the nosocomial infections^20^. Another recent report showed a higher frequency of occurrence of *bap* gene (56.6%) in *Staphylococcus* spp. (189 samples) isolated from bovine subclinical mastitis. Apart from this, frequency of *bap* gene occurrence was significantly higher in coagulase-negative strains as compared with coagulase-positive^21^. The involvement of polysaccharide-intercellular adhesin (*ica*-dependent) component of the *S. aureus* biofilm matrix has been studied comprehensively^22^. However, role of *ica*-independent mechanisms which is predominantly mediated by biofilm associated surface proteins (Bap,Aap, FnBPs etc.) in the stability of staphylococci biofilm matrix is poorly understood^13, 14, 23, 24^. Previous reports on *ica*-independent biofilm formation in *Staphylococci* showed a strong link between biofilm formation and cell wall associated proteins in particular, Bap,^16^ the accumulation associated protein (Aap)^24^ and a Bap-homologue protein (Bhp)^25^. A recent report shows that repeated domains contain an amyloidogenic peptide motif (-STVTVTF- derived from the C-repeat of the Bap), which is responsible for cell–cell interaction^26^. Therefore, Bap and Bap-like surface proteins could be an important target biofilm dispersal studies.

Dispersal mechanisms vary in different bacteria and this event is considered as a novel approach to treat drug resistant *S. aureus* which are common in body implants and catheter related infections^27^. Among the natural ways of *S. aureus* biofilm dispersal, Agr-mediated biofilm dispersal and secretion of major extracellular proteases, SspA, SspB, Aur and Scp as proenzymes were reported^28^. Theoretically, these enzymes may contribute to biofilm detachment but, very little is known about their role with regard to staphylococci. Recently an extracellular serine protease, Esp secreted by a subset of *Staphylococcus epidermidis*, shown to inhibit biofilm formation and nasal colonization by *S. aureus^29^*. Of late, it was shown that Esp has proteolytic activity specifically towards biofilm specific proteins that are associated with *S. aureus* biofilm formation and host-pathogen interaction^30^. Recent studies on *Staphylococci* chronic infections and biofilms as well as discovery of major dispersal mechanisms shifted the focus on development of dispersal-mediated treatment options for *S. aureus* biofilm infections^31^. With increasing number of reports on protein-based staphylococcal biofilms, it is speculated that protease based dispersion method would be highly effective^32^. In an earlier report, we showed that proteinase K can emulate the naturally produced proteases and can be used to enhance the biofilm dispersal through cleavage of surface proteins i.e. Bap-dependent *S. aureus* biofilm establishment^23^. In this additional report, we investigated whether this approach would be useful in general and has wider applicability by using five other *S. aureus* mastitis isolates. Apart from this we also investigated the effect of binding calcium to Bap on its stability against proteolytic activity of proteinase K. To address the problem, five *S. aureus* bovine mastitis isolates were included in the study along with *bap*–positive *S. aureus* V329 and a *bap*-isogenic mutant M556 as positive and negative controls respectively.

## Materials and Methods

### Microorganisms and culture conditions

A *bap*–positive *S. aureus* V329 and its isogenic mutant *S. aureus* M556 were used in this study along with five other mastitis isolates of *S. aureus* viz., SA7, SA10 SA33, SA252 and SA392. Bovine mastitis *S. aureus* strains SA7, SA10 and SA33 were procured from Karnataka Veterinary College, Bengaluru whereas SA352 and SA392 were procured from Madras Veterinary College, Chennai. All bovine mastitis *S. aureus* strains used in the study were isolated from infected site of bovine mastitis. M556 was generated by transposon insertion in the downstream part of *bap* gene of *S. aureus* V329 in such a way that Bap protein is synthesized but remains non-functional as cell wall anchoring region is truncated^16^. For each experiment, single colonies were picked from Tryptic Soy Agar (TSA) culture plates and inoculated in Tryptic Soy Broth (TSB) medium supplemented with 0.25% glucose (TSB-G) and incubated at 37oC at 150 rpm. Overnight grown cultures were used for all experiments after checking for culture purity. All experiments were performed at Biofouling and Biofilm Processes Section, Water and Steam Chemistry Division, BARC Facilities, Kalpakkam during the period of January 2013 to December 2014.

### Quantitative biofilm assay

Biofilm assay was performed in 96-well microtitre plates to estimate the inhibitory/dispersion action of proteinase K. The working concentration of proteinase K was chosen as 2 μg/mL in all the experiments. The overnight grown cultures of the *S. aureus* cells in TSB-G were diluted 1:40 in sterile TSB-G medium and added to the pre-sterilized 96 well flat bottom polystyrene microtitre plates. To estimate the inhibitory action *S. aureus* biofilms were grown in the presence of 2 μg/mL of proteinase K. To study dispersion, biofilms were grown on microtitre plates, washed after prescribed time and 200 μL of fresh TSB-G amended with 2 μg/mL of proteinase K was added to the wells and the plates were incubated at 37°C for 24 h. To study the effect of Ca^2+^ on proteolytic cleavage of Bap in terms of biofilm formation, V329 biofilms were grown at 37°C for 24 h in the presence of 2 μg/mL of proteinase K along with increasing concentration of Ca^2+^ in the range of 1.56 mM to 50 mM. V329 biofilm grown in the presence of only Ca^2+^ in the similar concentration range acted as control for Ca2+. After 24 h of incubation, biofilm growth was quantified. Biofilm quantification was done by classical crystal violet assay as described previously^3^. To dissolve the bound crystal violet, 33% acetic acid was used^33^. Biofilm growth was monitored in terms of absorbance at 570 nm using a multimode microplate reader (BioTek, USA).

### Planktonic growth studies

Overnight grown bacterial cultures were harvested and washed twice with phosphate buffered saline (PBS) and OD_600_ of each culture was set to 0.1. 100 μL of each resuspended culture was inoculated in 1900 μL of TSB-G and different concentrations of proteinase K. Cultures were incubated at 37°C and 150 rpm. Absorbance of each culture was recorded at different time intervals after vortexing for 5 sec, to re-suspend the settled cells.

### EPS extraction and quantification of biofilm matrix components

*Staphylococcus aureus* biofilm was grown for 48 h on glass slides immersed in 20 mL of TSB supplemented with 0.25% glucose (TSB-G). After 48 h, planktonic cells were aspirated and biofilm was gently washed twice with PBS. *S. aureus* biofilms were treated with 2 ug/mL proteinase K in TBS-G for 4 h at 37°C. After 4 h biofilm was gently washed with PBS and then remaining biofilm was scrapped and collected in 5 mL of PBS. Biofilm was disintegrated by gentle vortexing using glass beads. Five mL of the biofilm sample was centrifuged at 8000 rpm and 4oC for 30 min. Supernatant was collected and mixed with double volume of 90% chilled ethanol and kept at 4oC for overnight. EPS was collected by centrifugation at 10000 rpm and 4oC for 10 min. The supernatant was discarded and pellet was collected and dried at 60oC to remove ethanol. Pellet was resuspended in 100 μL PBS buffer. The protein and eDNA content in the resuspension was quantified using Qubit Fluorometer (Invitrogen, Carlsbad, CA, USA); the quantization protocol was followed as published by the vendor^34,35^. Glucose concentration as a measure of polysaccharide content was quantified by the method as described elsewhere^36^.

### Anti-biofilm activity of antibiotic-proteinase K treatment

To investigate the effect of proteinase K on the efficacy of antibiotics, proteinase K treatment was given a combination of gentamycin against biofilm forming *S. aureus* strains *viz* SA7, SA10, SA33 and SA352. The antibiotic concentrations were chosen as 10 and 50 times of the MIC (indicated as X and 5X respectively) against *S. aureus* planktonic cells. Proteinase K treatment was given in combination with X concentration of antibiotics. *S. aureus* biofilms were grown in microtitre plates at 37°C and 150 rpm. After 24 h, planktonic cells were aspirated by pipette and the biofilms were gently rinsed twice with sterile PBS. After rinsing, the biofilms were treated with antibiotics alone and antibiotics-proteinase K combinations. After 24 h, planktonic cells were aspirated and biofilms were gently rinsed twice with PBS. 200 μL of PBS was added to each well and the biofilm cells were dislodged by ultra-sonication for 5 min. Cells released from the biofilms were harvested and the viable cell count was obtained by plating on TSA media and incubated at 37°C overnight.

### Statistical Analysis

Two-tailed Student’s t test was used to determine the differences in biofilm formation between the groups. Differences were considered statistically significant when *P* value was < 0.05.

## Results

### Effect of proteinase K on biofilm development and planktonic growth of S. aureus strains

Figure 1 illustrates the results of biofilm inhibition assay, which shows that proteinase K treatment hampered the biofilm development of most *S. aureus* strains viz. SA7, SA10, SA33, SA352and *bap-* positive V329.All *S. aureus* strains except SA392 (weak biofilm producing strain) showed significant inhibition in biofilm growth when treated with 2 μg/mL. SA7, SA10, SA33, SA352 biofilms showed 84, 71, 83 and 68% reduction in biofilm growth in the presence of 2 μg/mL proteinase K. On the contrary, strains M556 and SA392 were found to be weak biofilm producers and there was no significant inhibition of biofilm formation in the presence of proteinase K. Planktonic growth studies of *bap-* positive V329 and *bap*-mutant M556 and other *S. aureus* strains was carried out in the presence of different concentration of proteinase K. Figure 2 and supplementary figure S1 show that there was no effect of proteinase K on the planktonic growth of either *S. aureus* strains when tested up to 32 μg/mL.

**Figure 1:**
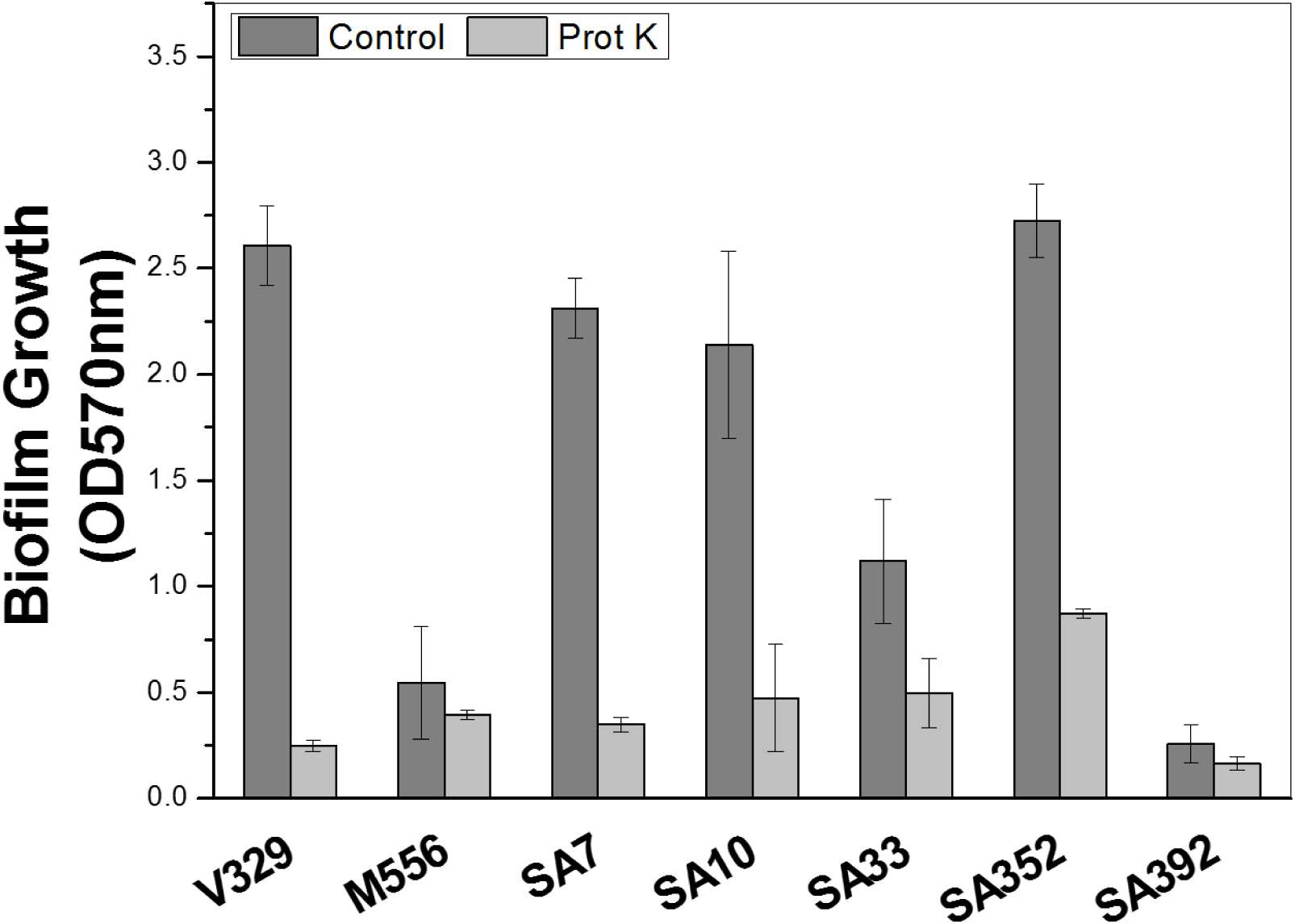
Effect of proteinase K on the growth of *S. aureus* biofilms.

**Figure 2:**
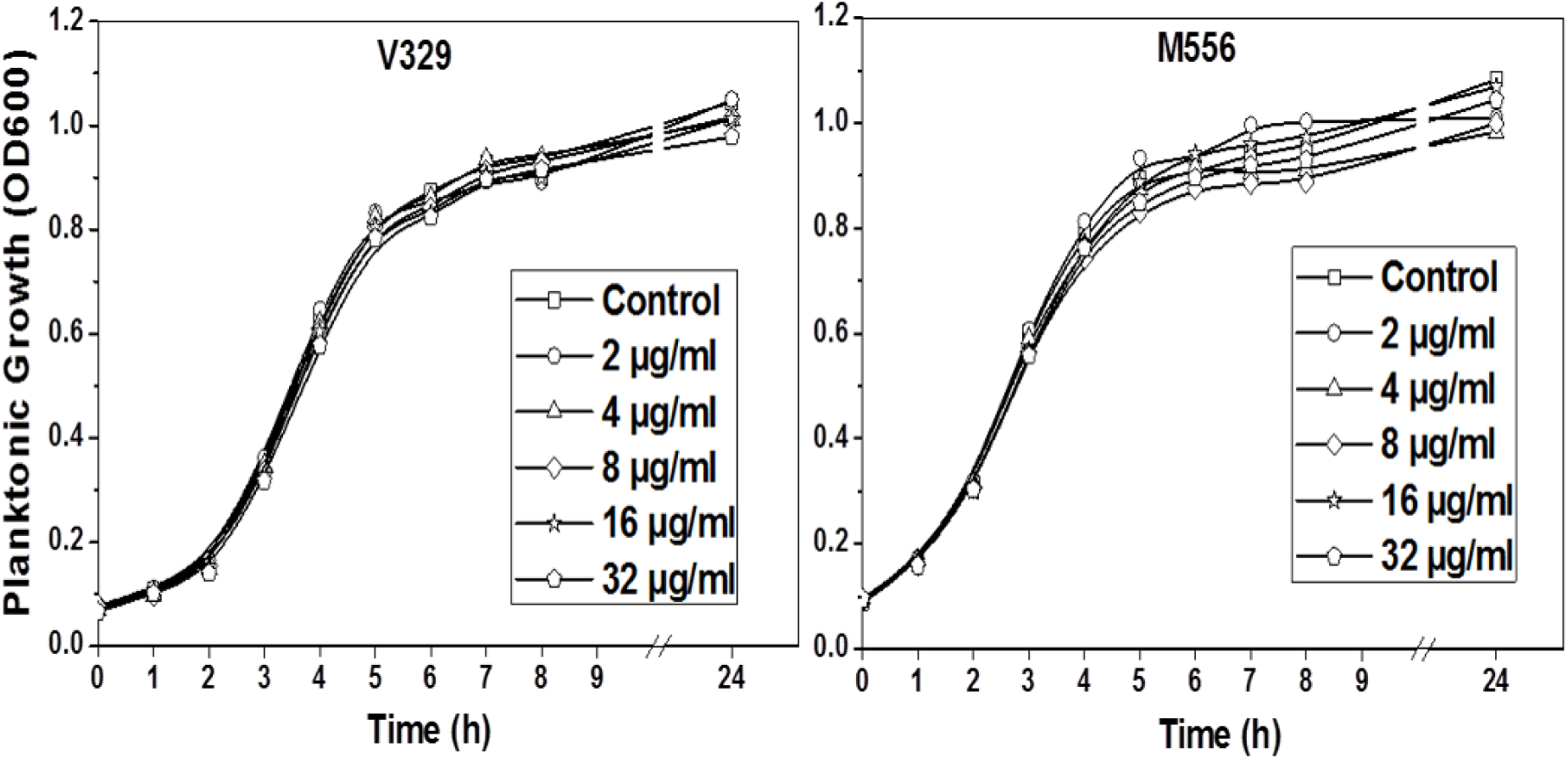
Effect of different concentrations of proteinase K on planktonic growth of *S. aureus* V329 and M556.

### Effect of different concentrations of Ca^2+^ on proteolytic degradation of Bap and biofilm development of V329

Biofilm assay in the presence of proteinase K with increasing concentrations of Ca^2+^ was carried out. Figure 3 shows that addition of increasing concentrations of Ca2^+^ had no effect on proteinase K mediated inhibition of biofilm development. In other words, Ca^2+^ did not affect the proteolytic degradation of surface protein Bap by proteinase K. On the other hand, lower concentration of Ca2^+^ (up to 6.25 mM) had no significant effect on V329 biofilm formation but higher concentrations showed an inhibitory effect which was found to be similar to the earlier report^23^.

**Figure 3:**
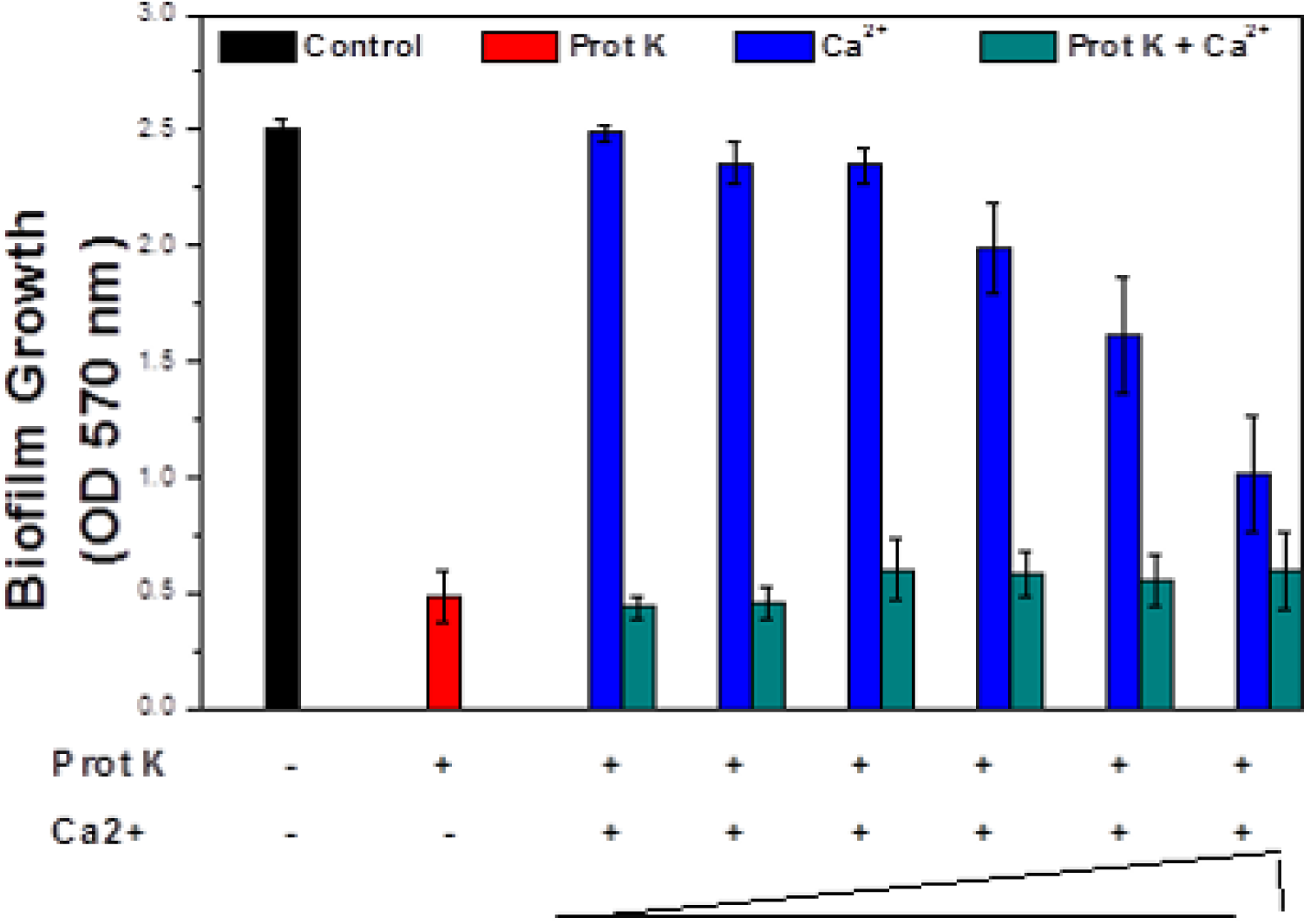
Effect of Ca^2+^ on proteinase K mediated biofilm inhibition in *bap*- positive *S. aureus* biofilm. Results are shown as mean ± 1 S.D

### Proteinase K enhances dispersal in S. aureus biofilms

In order to investigate the biofilm dispersal activity of proteinase K against *S. aureus* biofilms proteinase K treatment was given to 24 h old *S. aureus* biofilms. Figure 4 shows that proteinase K treatment of *S. aureus* biofilms caused a significant disruption of all *S. aureus* biofilms except M556 and SA392. Upon 2 μg/mL proteinase K treatment for 24 h, approximately 92.5 %, 23.4%, 90.1%, 95.8%, 81.9% and 60% biofilm dispersal was observed in V329, M556, SA7, SA10, SA33 and SA352 respectively. Though, proteinase K enhanced the biofilm dispersal, 100% biofilm removal could not be achieved in any case. Moreover, SA392 formed a weak biofilm as compared to V329 after 24 h, hence estimating SA392 biofilm dispersal was difficult and inappropriate, and therefore SA392 was not involved in further experiments.

**Figure 4:**
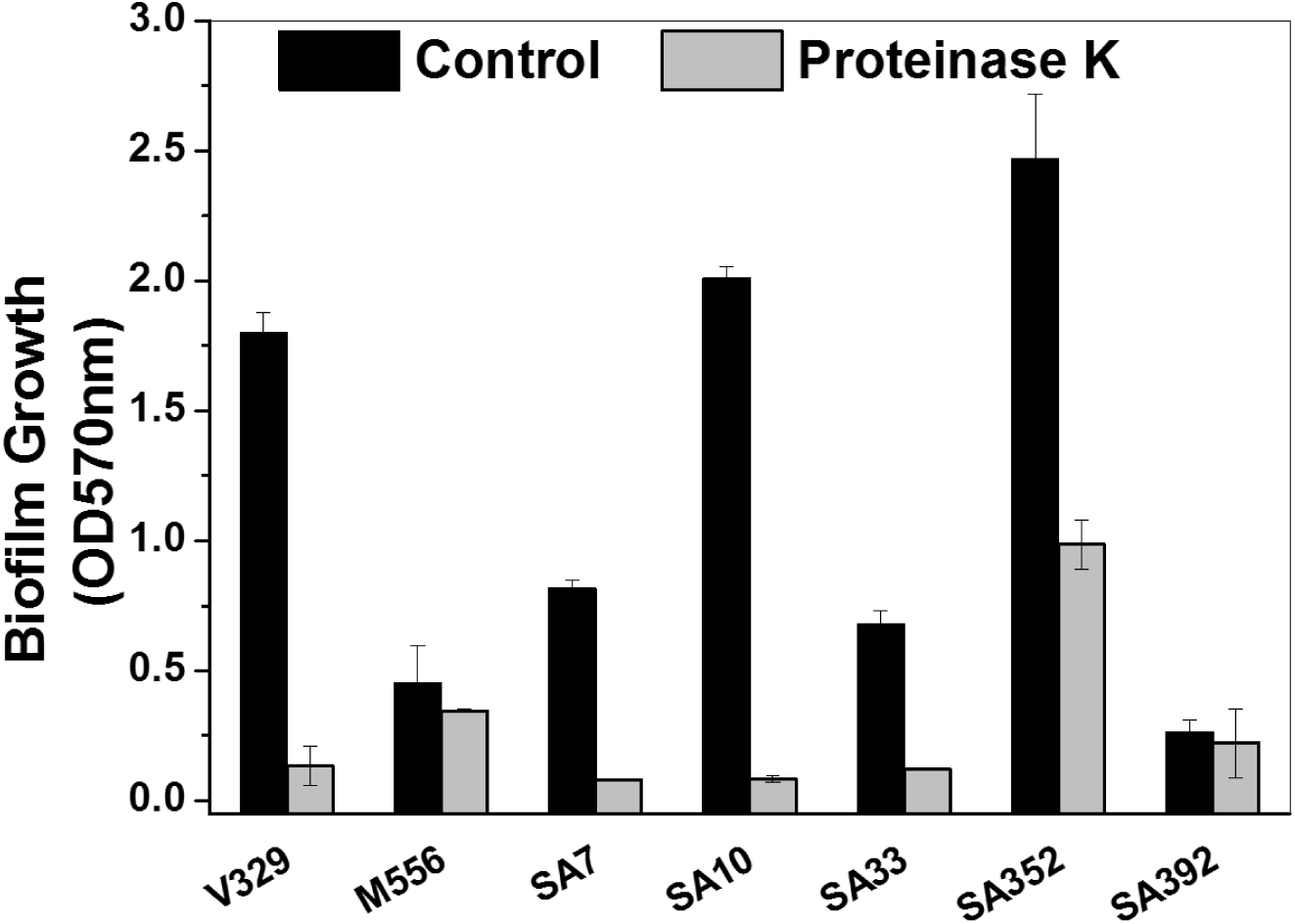
Dispersal of pre-grown 24 h old *S. aureus* biofilms by proteinase K. Results are shown as mean ± 1 S.D.

Figure 5 and supplementary table 1 show the constituents of *S. aureus* biofilms before and after the proteinase K treatment. *S. aureus* V329 biofilm had significantly higher amount of carbohydrate as well as eDNA (P < 0.05, n = 3) as compared to Bap mutant M556 biofilm. On the other hand M556 biofilm was comprised of significantly higher amount of biofilm matrix proteins (P < 0.05, n = 3). Figure 5 shows that there was significant decrease in the protein as well as eDNA content in V329 and M556 biofilms after proteinase K treatment, however there was no significant decrease in the carbohydrate content of biofilm matrix in either case.

**Figure 5:**
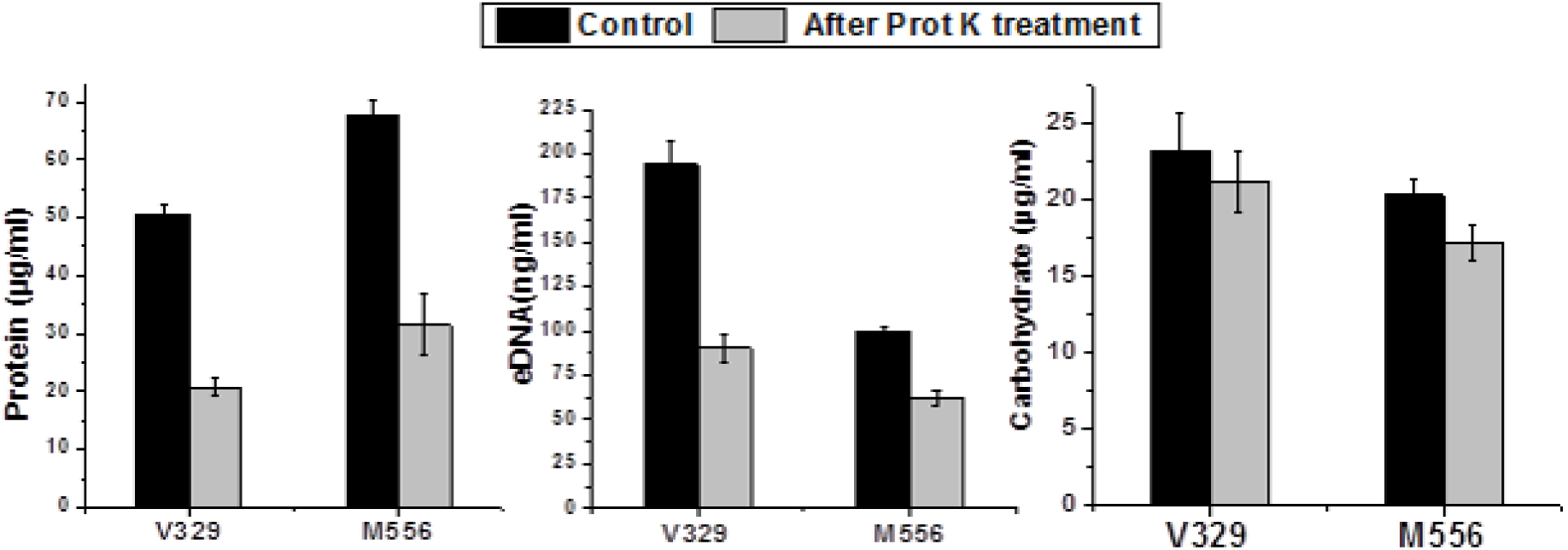
Constituents of *S. aureus* EPS before and after treatment of proteinase K. Results are shown as mean ± 1 S.D.

### Proteinase K enhances antibiotic efficacy against S. aureus biofilms

Figure 6 shows the reduction in cfu count in *S. aureus* biofilms (SA7, SA10, SA33 and SA352) upon treatment of gentamycin in various combinations. It was observed that addition of proteinase K in combination with gentamycin had more impact against *S. aureus* biofilm cells when compared to gentamycin alone. When the antibiotic concentration was increased by five times, there was no significant increase in log reduction of cfu count in any case. On the other hand, the addition of 2 μg/mL proteinase K in combination with antibiotics resulted in significant log reductions in each case. Proteinase K treatment caused an increase in the reduction in cfu in biofilms by 3.85, 5.0, 3.03 and 3.76 logs unit in the case of SA7, SA10, SA33 and SA352 respectively. On the other hand, only 0.45, 1.27, 0.88 and 0.79 log units enhancement was noticed when gentamycin concentration was increased 5 times.

**Figure 6:**
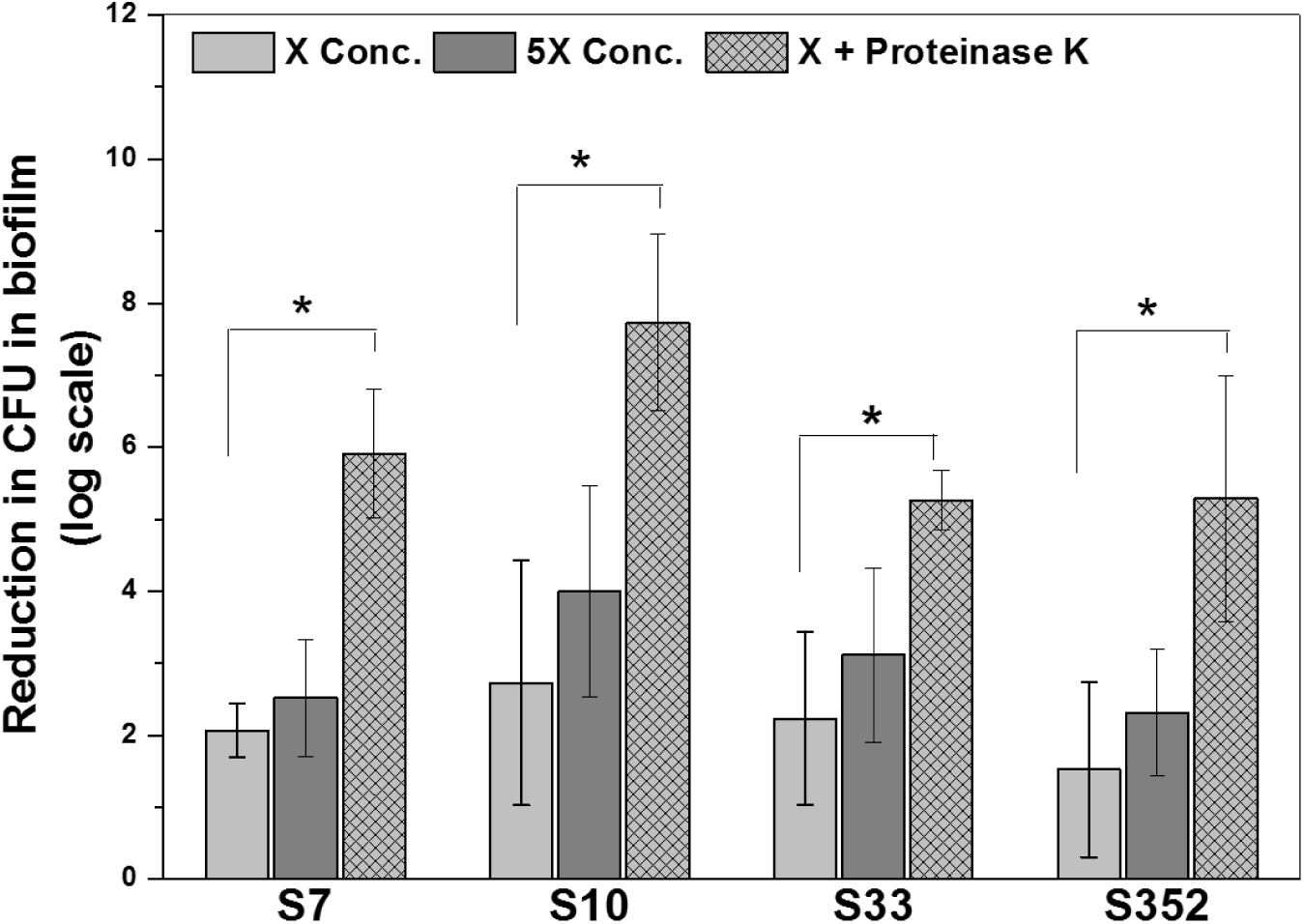
Antimicrobial efficacy of antibiotics in combination with Proteinase K against 24 h old *S. aureus* biofilms. Proteinase K was used at 2 μg/mL. Two concentrations of gentamycin were used; X = 150 μg/mL and 5X = 750 μg/mL. Asterisk shows significant difference when P < 0.05.

## Discussion

*Staphylococcus aureus* and *S. epidermidis* are the predominant microorganisms among HAIs such as catheters, cardiac valves and orthopaedic prostheses^6^. In clinical settings, *S. aureus* biofilms impose resistance to host immune/defence mechanisms and antimicrobial therapy thus enabling the bacterium to persist^37, 38^. During the last decade many reports have come up highlighting the important role played by surface proteins in early adhesion and establishment of biofilm^12^. It was anticipated that proteolytic cleavage of these biofilms would hamper the initial adhesion process and in turn progression of biofilm. The observations recorded in Figure 1 showed that proteinase K treatment had significantly impacted the biofilm development of most *S. aureus* isolates. In order to evaluate, whether inhibition of biofilm was due to the hampered growth or due to the life style switch over of the cells to planktonic form and effect of different concentration of proteinase k on planktonic cells were monitored. Figure 2 and supplementary Figure 1 show that there was no impact of proteinase K on cell viability, hence biofilm inhibition was due to proteolytic cleavage of surface proteins. These observations also reemphasize the important role played by Bap like surface proteins in biofilm development, as similar effect was not observed in *bap*- mutant M556.

In general, metal binding proteins are structurally more stable when they are bound to a specific metal and can show higher resistance towards proteolytic degradation^39^. As Bap contains four putative Ca^2+^ binding EF hand motifs, we evaluated whether binding of Ca^2+^ to Bap would confer any resistance against proteinase K mediated degradation and in turn biofilm dispersion. The result shown in Figure 3 indicates that there was no apparent difference between biofilms grown in the presence of proteinase K alone and proteinase K along with increasing concentrations of Ca^2+^. Whereas in another set of increasing concentrations of Ca^2+^ alone showed inhibition of V329 biofilm at higher concentrations as shown in an earlier report^23^. This result demonstrates that binding of Ca^2+^ ions to Bap had no effect on proteinase K mediated inhibition of V329 biofilm. In other words, biofilm assay using Ca^2+^ alone and Ca^2+^ with proteinase K showed that Ca^2+^ did not confer any immunity against proteolytic degradation of Bap.

As surface proteins also play an important role in stability of the *S. aureus* biofilms, preformed biofilms of various *S. aureus* strains (V329, M556, SA7, SA10, SA33, SA352 and SA392) were also tested with proteinase K. Figure 4 shows that proteinase K treatment also caused a significant dispersal of all pre-grown (24 h) *S. aureus* biofilms except M556 and SA392. Since in the case of M556, Bap protein does not remain attached to the cell wall, it remains non-functional and does not contribute in biofilm stability. M556 harbours functional *ica*-operon and hence could produce significant amount of biofilm after 48 h as shown in earlier report^16, 23^. Therefore, it can also be speculated that in weak biofilms by M556 and SA392, PIA might play a predominant role as their biofilms were not affected by proteinase K treatment.

The effect of proteinase K treatment on the composition of *bap* –positive and *bap*- mutant *S. aureus* biofilms was also investigated. After proteinase K treatment, a significant decrease in the protein and eDNA but not in the carbohydrate content in EPS was observed. eDNA is also known to play very important role in *S. aureus* biofilm stability^40^. As there was a significant decrease in eDNA content along with the biofilm matrix protein content, it is speculated that matrix proteins might also be involved in eDNA retention in the biofilm. Since Ca^2+^ binds with Bap^17^ as well as eDNA,^41^ it is speculated that Ca^2+^ might act as a cross-linking agent between Bap and eDNA thereby the presence of Bap can assist in retention of eDNA. Therefore, upon proteinase K treatment that degraded Bap, a significant amount of eDNA was also lost along with the proteins. As shown in Figure 5, M556 biofilm was comprised of significant amount of eDNA and carbohydrate, which suggest that in the absence of functional Bap protein, eDNA and carbohydrate i.e. polysaccharide polymers in matrix might play a crucial role in *S. aureus* biofilms. In M556 biofilm, matrix proteins do not contribute to biofilm stability despite having higher amount of protein content. The results obtained also indicate that matrix proteins were neither protected by sugars and DNA nor resistant to proteinase K and hence degraded by proteinase K. It is speculated that carbohydrate polymers retains the biofilm structure and do not allow the M556 biofilm to get dispersed upon proteinase K treatment.

It is an established fact that biofilm cells are extremely (1000 times or more) resistant to antibiotics as compared to planktonic cells due to physical as well as the genetic reasons^42, 43^. The proteinase K mediated dispersal of *S. aureus* biofilms suggested its potential use in enhancing the susceptibility of bacterial cells towards antibiotic treatment. Our earlier report showed that proteinase K treatment in combination with different antibiotics had a synergistic effect on efficacy of antibiotics^23^. On similar lines gentamycin efficacy in combination with proteinase K was investigated against four other biofilm forming *S. aureus* strains in this study. Result showed that proteinase K treatment significantly enhanced the efficacy of gentamycin against all *S. aureus* biofilm tested i.e., SA7, SA10 SA33 and SA352. As showed in our previous report proteinase K treatment increases the surface to volume ratio and roughness coefficient of biofilm. Thus, the enhanced values of surface to biovolume ratio and roughness coefficient after proteinase K treatments leave more biofilm surface available for antibiotics action. Moreover, it was also shown that proteinase K treatment significantly decreased the average diffusion distance as well as maximum diffusion distance^23^. This enhances the antibiotics penetration in biofilms and in turn its efficacy. Thus, proteinase K shows synergistic effect when associated with antibiotics for biofilm removal.

In the recent past few years, enzyme based *S. aureus* biofilm disruption has emerged as a promising strategy to combat biofilm-related persistent infections as enzyme based antibiotic treatment enhances the antibiotic sensitivity of microbial biofilm^31, 44^. Among such enzymes DNase I, Dispersin B and proteinase K are commercially produced. Dispersal mechanisms using such enzymes could be utilized in the prevention of biofilm formation associated with implanted medical devices^31^. Several studies have found that pretreatment of polymeric surfaces^45^ or local delivery of dispersal agents from the implanted device^10^ should prevent biofilm development. Few studies have been done with recombinant human DNase I to reduce the antigenicity of DNase I enzyme and has been shown promising results in terms of antibiofilm activity against *S. aureus* biofilm^46^. While these approaches sound promising, there are several concerns need to be thoroughly addressed before clinical trials. For example, if the antibiotic treatment along with the proteinase K fails to fully eradicate the dispersed microbial cells, it might result in acute infections. Thus, more studies need to be performed to confirm dispersal mechanisms in relevant animal models of infection before treating *S. aureus* biofilm infections in clinical set ups.

In conclusion, the potential application of this study denotes that, proteinase K mediated biofilm removal has a wider applicability. Proteinase K can be used in biofilm dispersion as well as for anti-biofilm activity as most of the biofilms do comprise of surface proteins as a major constituent of biofilm matrix. The present study show a wider applicability of proteinase K treatment against *S. aureus* biofilms and it also establishes the fact that antibiotics in combination with proteinase K can be more effective in controlling *S. aureus* biofilm mediated infection. Apart from clinical biofilms, proteinase K has also been shown to impair *Listeria monocytogenes* biofilm formation and induce dispersal of pre-existing biofilms in food industry^44^. This study shows that targeting biofilm matrix components by specific enzymes, e.g. surface proteins could be a potential approach towards controlling biofilms related problems.

## Acknowledgements

### Conflict of interest

Authors of the manuscript declare no conflict of interest.

